# Short communication: Accounting for nuclear and mito genome in dairy cattle breeding - a simulation study

**DOI:** 10.1101/2023.11.20.567907

**Authors:** Gabriela Mafra Fortuna, B.J. Zumbach, M. Johnsson, I. Pocrnic, G. Gorjanc

**Author notes:** Corresponding author: Gabriela Mafra Fortuna, The Roslin Institute, The University of Edinburgh, Easter Bush Campus, Midlothian EH25 9RG.

## Abstract

Mitochondria play a significant role in numerous cellular processes through proteins encoded by both nuclear genome (nDNA) and mito genome (mDNA). While the variation in nDNA is influenced by mutations and recombination of parental genomes, the variation in mDNA is solely driven by mutations. In addition, mDNA is inherited in a haploid form, from the dam. Cattle populations show significant variation in mDNA between and within breeds. Past research suggests that variation in mDNA accounts for 1-5% of the phenotypic variation in dairy traits. Here we simulated a dairy cattle breeding program to assess the impact of accounting for mDNA variation in pedigree-based and genome-based genetic evaluations on the accuracy of estimated breeding values for mDNA and nDNA components. We also examined the impact of alternative definitions of breeding values on genetic gain, including nDNA and mDNA components that both impact phenotype expression, but mDNA is inherited only maternally. We found that accounting for mDNA variation increased accuracy between +0.01 and +0.05 for different categories of animals, especially for young bulls (+0.05) and females without genotype data (between +0.01 and +0.03). Different scenarios of modelling and breeding value definition impacted genetic gain. The standard approach of ignoring mDNA variation achieved competitive genetic gain. Modelling, but not selecting on mDNA expectedly reduced genetic gain, while optimal use of mDNA variation recovered the genetic gain.

---

Most breeding research and applications focus on how variation between and within nuclear genomes (nDNA) affects economically important traits. However, other genomic elements, such as mitochondrial genomes (mDNA), may also impact response to selection (Bell et al., 1985). In dairy cattle, ∼30% of the phenotypic variation for milk yield is associated with variation in the nDNA (e.g. García-Ruiz et al., 2016), which is a molecule of ∼3Gb (e.g., Srirattana and St. John, 2017). In contrast, variation in the mDNA, which spans only ∼16kbp (e.g., Srirattana and St. John, 2017), is associated with 1-5% of the phenotypic variation (Bell et al., 1985; Brajkovic et al., 2023; Schutz et al., 1992). These proportions indicate that dairy breeding could benefit from accounting for mDNA variation. With the advances in SNP array technologies and the increasing accessibility of whole-genome sequencing, it will soon be possible to routinely include the variation in nDNA and mDNA in genetic evaluations and practical breeding programs. Mitochondria, have critical roles in cellular processes. They generate energy, synthesize adenosine triphosphate, and contribute to metabolic homeostasis etc. Their evolution from an autonomous prokaryote involved gene loss and transfer, leading to close integration with the host’s nDNA (Ladoukakis and Zouros, 2017). Unlike nDNA, mDNA is transmitted between generations via maternal lineages without recombination (e.g., Roger et al., 2017; Sato and Sato, 2013). This mechanism is thought to avert conflicts and safeguard the genome from selfish genes and the consequential reduction in fitness (Hastings, 1992). Research indicates a higher mutation rate in mDNA compared to nDNA, which may result from the intra-mitochondrial environment (Ladoukakis and Zouros, 2017). Mitochondrial DNA is categorized into haplogroups, reflecting inter-population variation. However, variation within populations (breeds) is also observed (Dorji et al., 2021). This diversity is associated with various traits. In humans, mDNA polymorphisms are associated with genetic disorders and variations in quantitative traits (Stewart & Chinnery, 2015). In dairy cows, mDNA plays a role in milk yield and composition (Bell et al., 1985; Schutz et al., 1992), possibly due to the energy-intensive lactation process. All this suggests that mDNA variation should be accounted for in breeding programs. Gibson *et al*. (1997) demonstrated that even small contributions from mDNA to the total variation of a trait can drive differences in performance between maternal lineages. The maternal inheritance of mDNA enables tracking maternal lineages from founders via pedigrees (Schutz et al., 1992). Spehar et al. (2017) estimated that variation between pedigree maternal lineages accounted for 2-3% of the phenotypic variance for milk yield in the Croatian Holstein population, while a genomic analysis in the same population accounted for up to 5% of the phenotypic variance (Brajkovic et al., 2023). An earlier simulation study accounting for variation between maternal lineages suggested an increase in the accuracy of estimated breeding values for cows by 0.01 when the maternal lineages accounted for 2.5% of the phenotypic variance and 0.04 when accounted for 10% (Boettcher et al., 1996). Gibson *et al*. (1997) also highlighted that accounting for maternal lineages reduces the bias of estimated breeding values for cows because they transmit both nDNA and mDNA components of their breeding value to their offspring. Considering the limited number of studies and the ongoing questions, this work aims to expand the past research by evaluating (i) the impact of accounting for the mDNA variation on the accuracy of genetic evaluation and (ii) the impact of alternative breeding value definitions (including nDNA and mDNA components) on genetic gain in a simulated dairy cattle breeding program.

## Materials & Methods

A dairy cattle breeding scheme was simulated with the R package AlphaSimR (Gaynor *et al*., 2020). All the simulation scripts are available at https://github.com/HighlanderLab/gfortuna_mtdna_breed. We simulated nDNA and mDNA independently. To simulate nDNA chromosome haplotypes, we used the coalescent simulator MaCS (Chen et al., 2009) as implemented in AlphaSimR with the “CATTLE” parameters for

10 diploid chromosomes (to reduce computation time) with 10^8^ base pairs each, mutation rate of 2.5x10^−8^, recombination rate of 1x10^−8^, and historical effective population sizes as described in MacLeod *et al*. (2013). We chose 1,000 loci per chromosome as SNP markers and another 1,000 as QTL. For the mDNA haplotypes, we considered 1 haploid chromosome with 16,202 base pairs, mutation rate of 2.5x10^−7^, and no recombination. We set mDNA effective population size to 1,000 in the most recent generation. We increased the historical effective population sizes from MacLeod *et al*. (2013) (see GitHub) to obtain over 1,000 polymorphic loci in line with literature (Brajkovic et al., 2023; Dorji *et al*., 2021). We chose all polymorphic loci (on average 1,084 across 10 replicates) in the mDNA as SNP markers. We evaluated a scenario where all or only one randomly chosen SNP were/was a QTL. Both simulated nDNA and mDNA haplotypes were randomly allocated to nuclear and mito genomes of 1,000 founding individuals. The nDNA was then passed between generations with recombination in diploid form, while mDNA was passed only from mothers to their progeny without recombination in haploid form. We defined one polygenic trait with a heritability of 0.3, partitioned between nDNA 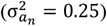 and mDNA 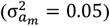 components in the base population. QTL allele substitution effects for nDNA and mDNA were sampled from a Gaussian distribution that generated targeted genetic variances and heritability after environmental variation from Gaussian distribution was added. The trait was expressed only in cows and was generated as:

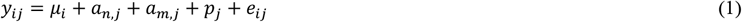

where *y*_*ij*_ is the phenotype of animal *j* in lactation *i, µ*_*i*_ is the population mean for lactation *i, a*_*n,j*_ is the nDNA breeding value for animal *j* (nTBV), *a*_*m,j*_ is the mDNA breeding value for animal *j* (mTBV) (which was the same as for its mother, maternal grandmother, etc.), *p*_*j*_ is the permanent environment effect of animal *j* sampled from 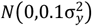, and *e*_*ij*_ is the environmental effect sampled from 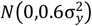. Therefore, in the base population the simulated phenotypic variance was 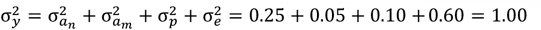.

We defined an individual *j*’s breeding value in two ways: (i) as nDNA breeding value (nTBV) and (ii) as the sum of nDNA and mDNA breeding values (tTBV = nTBV + mTBV). The definition (ii) is correct for phenotype expression in males and females because both sexes have nDNA and mDNA, and for inheritance in females because only they transmit nDNA and mDNA to the next generation. Hence, for females, definition (ii) should always be used, while for males, definition (i) should be used for selection and definition (ii) should be used for modelling their own phenotype data. However, male phenotypes are seldom modelled in dairy breeding.

We analyzed the phenotype data to estimate breeding values (EBV) using the generative model (1) with pedigree- and genome-based information and accounting for the mDNA variation or not. In the pedigree-based model, we assumed 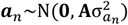 with **A** being the pedigree relationship matrix for the nuclear genome of dimension equal to the number of animals in the pedigree, and 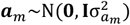 with **I** being an identity matrix of dimension equal to the number of distinct maternal founder lineages. In the genome-based model, we assumed 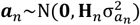 with **H**_*n*_ being the “single-step” joint pedigree- and genome-based relationship matrix for nDNA with dimension equal to the number of animals in the pedigree (Aguilar et al., 2010), and 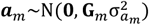 with **G**_*m*_ being the genome-based relationship matrix for mDNA with dimension equal to the number of distinct mDNA in the data, *n*_*m*_. Note that *n*_*m*_ was smaller than the number of distinct maternal founder lineages in pedigree. We calculated the mDNA genomic relationship matrix as **G**_*m*_ = **M**_**m**_**M**_**m**_^T^/*k*, where **M**_**m**_ = **W**_*m*_ – **P**_*m*_ with **W**_*m*_ an *n*_*m*_ × *n*_*s*_ matrix of 0s (for ancestral allele) and 1s (for mutation) for *n*_*m*_ distinct mDNA and *n*_*s*_ polymorphic loci (we assume we know all these loci by sequencing mDNA within pedigrees; Brajkovic et al., 2023), **P**_*m*_ a matrix of mutation frequencies, *k* = 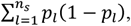, and *p*_*l*_ the mutation frequency at locus *l*. To speed up simulations, we fixed variance components to simulated values during the burn-in phase (see the next paragraphs). After the burn-in phase, we estimated the variance components and used these new estimates for the remainder of the simulation. We fitted all the models with the BLUPF90 suite (Misztal et al., 2018).

We evaluated sixteen scenarios driven by three factors across 10 replicates. All scenarios included a burn-in step of 10 years of progeny testing-based selection. The first factor was the breeding scheme: (i) progeny testing-based selection (PT) and (ii) genomic selection (GS). We simulated a 20-year breeding program considering overlapping generations, generating 35,179 animals annually. This population size was determined to produce 5 Elite Sires every year with a selection intensity of 0.9. In the PT scenarios, the male selection pathway had a generation interval of 6 years, with 4 years for the progeny test. This time was reduced to 2 years in the GS scenarios. To ensure accurate genetic evaluations, each Waiting Bull was required to have at least 100 phenotyped daughters at the time of testing. The female selection pathway involved 5 lactations over a 7-year generation interval. We categorized the animals into six groups: Elite Dams (top 250 first-lactation cows), Commercial (best 70% first-lactation cows after Elite Dams selection), Heifers (7,110 females without lactation record), Young Bulls (97% male offspring from Elite categories – nucleus population), Waiting Bulls (top 50 Young Bulls based on breeding values), and Elite Sires (5 highest-rated males based on breeding values). Elite Dams and Commercial females were replaced at a 30% rate/year and 100% after their 5^th^ lactation, while all Elite Sires were replaced after 5 years in the category. In GS scenarios, 10,200 genotypes were generated each year. The reference population began with 8,944 genotyped females. New genotypes, 613 males and 2,461 females, were added every year. All genotyped animals were from the nucleus. Old records were removed to keep the number of genotyped animals within the 25,000 limit of the free-version of BLUPF90.

The second factor was a statistical model accounting for mDNA variation or not and selection on different definitions of breeding value: (a) using a standard model (without mDNA) and selecting both females and males on their nDNA EBV (STANDARD), (b) using a model with mDNA and selecting both females and males on their nDNA EBV (BASELINE), (c) using a model with mDNA and selecting females on their nDNA plus mDNA EBV and selecting males on their nDNA EBV (OPTIMUM), and (d) using a model with mDNA and selecting both females and males on their nDNA plus mDNA EBV (EXTREME). We consider the OPTIMUM scenario to be the correct strategy to be used, hence the name.

The third factor was the assumption that all or just one polymorphic locus in mDNA is a QTL.

We evaluated all the scenarios with (i) accuracy of the last year of genetic evaluation as the correlation between true and estimated breeding value for scenarios STANDARD and BASELINE - because accuracy does not change with the OPTIMUM and EXTREME scenarios and (ii) genetic gain for mTBV, nTBV, and tTBV = mTBV + mTBV (in units of tTBV standard deviation from year 10) as the mean after 20 years of selection for scenarios STANDARD, BASELINE, OPTIMUM, and EXTREME.

## Results

We present results separately for the following five categories: (1) Heifers, (2) 1^st^ lactation cows (Cows1), (3) cows with 2 to 5 lactation records (Cows2-5), (4) young bulls, males without progeny; and (5) proven bulls, progeny tested males. For the GS scenarios, the female categories were split to show the difference between genotyped (g) and non-genotyped (ng) animals. We present only the scenario where all mDNA polymorphic loci were QTL (the other scenario with a single QTL was qualitatively the same).

In both PT and GS cases, accounting for mDNA variation increased the accuracy of nDNA EBV between 0.01 and 0.05, depending on the animal category (Table 1). In the PT case, the highest increase was observed for the young bulls (+0.05) and Cows2-5 (+0.03). The same was observed in the GS, accuracy of nDNA increased for 0.05 in young bulls and 0.03 for non-genotyped Cows2-5, while for genotyped Cows2-5 the increase was 0.02. The accuracy of mDNA EBV in both PT and GS was close to one for all animal categories (results not shown), which is expected given the lack of recombination in mDNA. Variation between simulation replicates was substantial.

**Table 1.** Accuracy (average ± standard deviation across replicates) of estimated breeding values for different animal categories in the model with nDNA breeding value (STANDARD) or with nDNA and mDNA breeding value (BASELINE) with the pedigree-based model in progeny testing-based selection (PT) or with genome-based model in genomic selection (GS) scenarios.

| Scenario | Category | STANDARD | BASELINE | Difference |
| --- | --- | --- | --- | --- |
| PT | Heifers | 0.47 ± 0.01 | 0.48 ± 0.01 | 0.01 ± 0.01 |
| | Cows1 | 0.61 ± 0.01 | 0.63 ± 0.00 | 0.02 ± 0.01 |
| | Cows2-5 | 0.56 ± 0.01 | 0.58 ± 0.01 | 0.03 ± 0.01 |
| | Young bulls | 0.36 ± 0.03 | 0.41 ± 0.05 | 0.05 ± 0.06 |
| | Proven bulls | 0.74 ± 0.02 | 0.73 ± 0.02 | 0.02 ± 0.03 |
| GS | Heifers (ng) | 0.41 ± 0.01 | 0.42 ± 0.01 | 0.01 ± 0.01 |
| | Heifers (g) | 0.73 ± 0.01 | 0.74 ± 0.01 | 0.01 ± 0.01 |
| | Cows1 (ng) | 0.56 ± 0.00 | 0.58 ± 0.00 | 0.02 ± 0.01 |
| | Cows1 (g) | 0.77 ± 0.00 | 0.78 ± 0.00 | 0.01 ± 0.01 |
| | Cows2-5 (ng) | 0.52 ± 0.01 | 0.56 ± 0.00 | 0.03 ± 0.01 |
| | Cows2-5 (g) | 0.73 ± 0.01 | 0.75 ± 0.01 | 0.02 ± 0.01 |
| | Young bulls (g) | 0.71 ± 0.02 | 0.76 ± 0.01 | 0.05 ± 0.02 |
| | Proven bulls (g) | 0.78 ± 0.01 | 0.78 ± 0.02 | 0.00 ± 0.02 |

g – genotyped; ng – non-genotyped

Genetic gain for mTBV, nTBV, and tTBV after 20-years of breeding with different estimation and selection scenarios is shown in Figure 1. Modelling but not selecting on mDNA variation (BASELINE) reduced genetic gain for tTBV compared to not modelling it (STANDARD). This was due to the lack of genetic gain for mTBV with the BASELINE scenario, while the STANDARD scenario partially captured the mTBV variation even without direct modelling and selection – via “mTBV-biased” estimation of nTBV. Modelling and selecting on mDNA variation recovered genetic gain for tTBV in OPTIMUM and EXTREME scenarios. There was an indication of increased gain for mTBV with the OPTIMUM and EXTREME scenarios, though variation between replicates was substantial, particularly for the EXTREME scenario. These trends were similar for the PT and GS breeding scheme.

**Figure 1.**
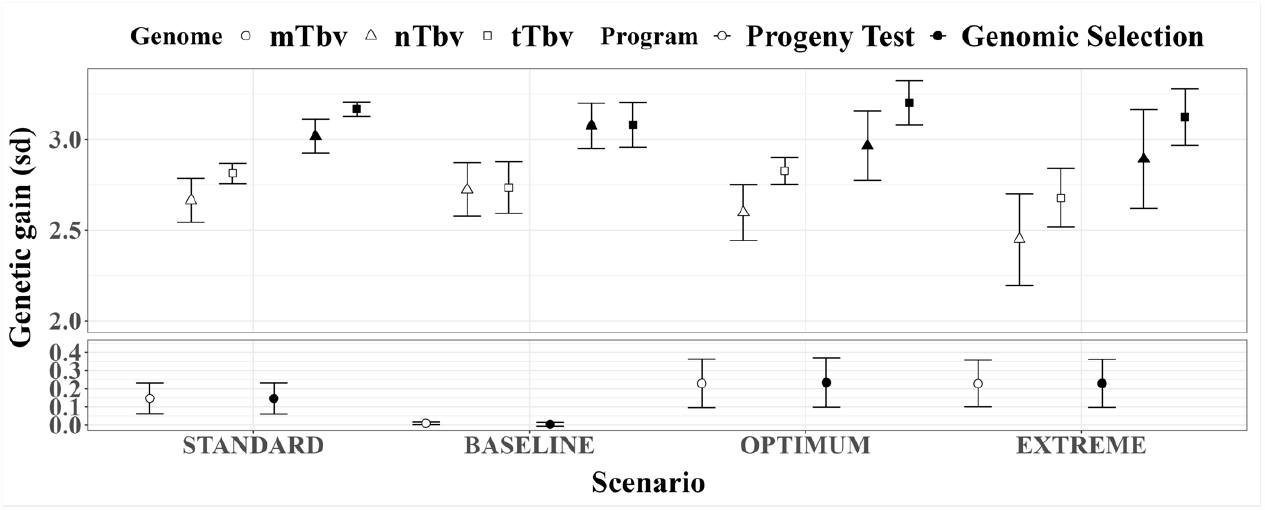
Standardized genetic gain (average ± standard deviation across replicates) after 20 generations of selection. Circles indicate results for mito true breeding values (mTBV), triangles for nuclear (nTbv) and squares for total true breeding value (nTBV + mTBV) - the definition of breeding value in estimation and selection varied according to simulation scenario (see Methods). Empty symbols indicate results with progeny testing-based selection and full symbols indicate results with genomic selection.

## Discussion

This study shows that considering mDNA variation can improve dairy breeding by increasing accuracy of selection, but genetic gain depends on how mDNA variation is modelled and selected upon. In the following, we discuss: 1) mDNA causal sites, 2) mDNA demography, 3) mDNA heteroplasmy, and 4) interactions between mDNA and nDNA.

### mDNA causal sites

We simulated scenarios with one mDNA segregating site as QTL as well as all segregating sites as QTL, which made no qualitative difference to results. When simulating one QTL, we assumed this site was the source of all observed phenotypic variation due to mDNA. However, when considering all sites, the effect of individual QTL was reduced. The absence of recombination creates strong linkage between loci along the mDNA, making the sum of several small-effect QTLs effectively equal to that of a single large-effect QTL. This also explains why the mDNA EBV had almost perfect accuracy, which in turn explains differences between the tested scenarios for modelling and selecting on mDNA variation. Future studies with collected data will give more information about the QTL, realized accuracies, and genetic gain.

### The impact of mDNA demography

Our initial simulations of mDNA, indicated that we need larger N_e_ for mDNA than for nDNA (despite an order of magnitude larger mutation rate in mDNA) to obtain a number of segregating sites in mDNA in line with the observed variation both within and across cattle populations (Dorji et al. 2021; Brajkovic et al., 2023;). As expected, we noticed that the lower the diversity among the mDNA, the smaller the impact of accounting for mDNA in breeding value estimation, in line with Boettcher et al. (1996). Theoretically, effective population size (N_e_) for mDNA equals the number of females or a quarter of the nDNA N_e_, though these relationships depend on the sex ratio (Birky et al., 1983). In a recent study, Cubric-Curik et al. (2021) inferred the demographic trend in N_e_ for mDNA in cattle. They found that mDNA N_e_ is increasing over time, which is opposite to results for nDNA in dairy cattle (MacLeod *et al*., 2013), but in line with the large diversity observed in mDNA (∼1,000 polymorphic sites out of 16,202 base pairs). The limited effect of accounting for mDNA observed in this study could be an underestimation due to the mismatched demographic parameters for the mDNA in our simulation. More research is needed to estimate demographic trends in nDNA and mDNA jointly and to understand how inheritance of these two DNA molecules interplays with modelling and selection.

### Heteroplasmy

We did not consider the presence of multiple mDNA copies with possible differences in mitochondria, known as heteroplasmy (Stewart & Chinnery, 2015). Future research should consider this additional variation, which is challenging since heteroplasmy varies between cells, tissues, and time points.

### nDNA-mDNA interaction

We did not consider interactions between mDNA and nDNA. Studies suggest that incompatibility between the two DNA can lead to mitochondrial malfunction, decreasing energy production efficiency and increasing oxidative damage (Pozzi *et al*., 2022, Ward *et al*., 2022). This factor could be relevant, and therefore, additional research is required, especially in the context of taurine-indicine crossbreeding systems (Ward *et al*., 2022).

**In conclusion**, the results show that accounting for mDNA variation can impact the success of dairy breeding. This study provides a genomic update of Boettcher et al. (1996) with additional results. With the increasing relevance of selection among females, especially for use in egg-transfer schemes, accounting for the mDNA will improve the accuracy of selecting more productive cows.

### Notes

GMF acknowledges the support from the Erasmus+ Program of the European Union to the European Masters in Animal Breeding and Genetics (EMABG).

MJ acknowledges support from Formas — a Swedish Research Council for Sustainable Development (Dnr. 2020-01637).

IP and GG acknowledge support from the BBSRC to The Roslin Institute (BBS/E/D/30002275, BBS/E/RL/230001A, and BBS/E/RL/230001C) and The University of Edinburgh.

For open access, the authors have applied a CC BY public copyright license to any Author Accepted Manuscript version arising from this submission.

No animal subjects were used in the study; all data was obtained from computer simulations. The authors have not stated any conflicts of interest.

